# Neuronal Adaptation to the Value Range in the Macaque Orbitofrontal Cortex

**DOI:** 10.1101/399071

**Authors:** Katherine E. Conen, Camillo Padoa-Schioppa

## Abstract

Economic choice involves computing and comparing the subjective values of different options. The magnitude of these values can vary immensely in different situations. To compensate for this variability, decision-making neural circuits adapt to the current behavioral context. In orbitofrontal cortex (OFC), neurons encode the subjective value of offered and chosen goods in a quasi-linear way. Previous work found that the gain of the encoding is lower when the value range is wider. However, previous studies did not disambiguate between neurons adapting to the value range or to the maximum value. Furthermore, they did not examine changes in baseline activity. Here we investigated how neurons in the macaque OFC adapt to changes in the value distribution. We found that neurons adapt to both the maximum and the minimum value, but only partially. Concurrently, the baseline response is higher when the minimum value is larger. Using a simulated decision circuit, we showed that higher baseline activity increases choice variability, and thus lowers the expected payoff in high value contexts.

## Introduction

Neuronal adaptation takes place throughout the brain. While its function is not fully understood, in sensory systems adaptation may contribute to homeostatic regulation (Benucci, Saleem, & Carandini, 2013; Hengen, Lambo, Van Hooser, Katz, & Turrigiano, 2013), efficient perceptual representation (Adibi, McDonald, Clifford, & Arabzadeh, 2013; Dan, Atick, & Reid, 1996; Gutnisky & Dragoi, 2008; Lewicki, 2002), and sharper behavioral performance (Krekelberg, van Wezel, & Albright, 2006; Liu, Macellaio, & Osborne, 2016). Context adaptation has also been observed in the neuronal representation of subjective values. Studies in non-human primates found adaptive coding in several brain regions, including orbitofrontal cortex (OFC) (Kobayashi, Pinto de Carvalho, & Schultz, 2010; Padoa-Schioppa, 2009; Yamada, Louie, Tymula, & Glimcher, 2018), anterior cingulate cortex (Cai & Padoa-Schioppa, 2014), and the amygdala (Bermudez & Schultz, 2010; Saez, Saez, Paton, Lau, & Salzman, 2017). In humans, experiments measuring BOLD activity have shown context adapting value signals in ventromedial prefrontal cortex (vmPFC), ventral striatum, and other brain areas (Burke, Baddeley, Tobler, & Schultz, 2016; Cox & Kable, 2014; Elliott, Agnew, & Deakin, 2008). More recent work has begun to explore the behavioral implications of value adaptation using a combination of experimental and theoretical approaches. One study found that adaptation in OFC reduces variability in value-based decisions, increasing the subject’s expected payoff (Rustichini, Conen, Cai, & Padoa-Schioppa, 2017). Other work suggests that value adaptation on a shorter time scale may produce irrational decision patterns (Soltani, De Martino, & Camerer, 2012; Yamada et al., 2018).

Despite this growing interest, our understanding of value adaptation is incomplete. In particular, previous studies did not clearly distinguish between neurons adapting to the range of values and neurons adapting to the maximum value available in a given context (Cox & Kable, 2014; Kobayashi et al., 2010; Padoa-Schioppa, 2009). Furthermore, these studies focused exclusively on the gain of value encoding (Cox & Kable, 2014; Kobayashi et al., 2010; Padoa-Schioppa, 2009; Rustichini et al., 2017) and did not examine potential changes in the overall response (i.e., changes in offset). In this study, we developed a task that allowed us to address these issues. We focused on the OFC, an area engaged in value-based decisions (Fellows, 2011; Padoa-Schioppa & Conen, 2017; Rudebeck & Murray, 2014; Schultz, 2015; Wallis, 2012).

We examined how value-encoding cells adapt to changes in both the maximum and the minimum of the value distribution. Neurons adapted to both maximum and minimum values, but responses did not remap completely to the new value range. Importantly, partial remapping reflected the final adapted state of neurons, not simply an incomplete temporal process. One byproduct of partial adaptation was an increase in the baseline response in contexts with a higher minimum value. Simulating a linear decision network, we showed that this change in baseline activity could increase choice variability, reducing the subject’s overall payoff. However, this theoretical loss is minor compared to the effect of narrowing the dynamic range. Incomplete adaptation may allow the circuit to maintain information about the overall value of the context, at the cost of a slight decrease in expected payoff.

## Results

To measure neuronal adaptation, we trained animals to perform a modified version of a juice choice task (Fig.1A). The task consisted of 2-3 blocks of ∼ 250 trials. Within each block, the monkey chose between two juices labeled A and B (with A preferred). The quantity of juice offered varied pseudo-randomly within a set range, defined by a minimum and maximum value (*V*_*min*_ and *V*_*max*_). In a given block, each juice could be offered in a “high”, “low”, or “wide” range (Fig.1B). Between blocks, the range of offers for each juice changed in one of six ways: *V*_*max*_ increased / decreased, *V*_*min*_ increased / decreased, or both *V*_*max*_ and *V*_*min*_ increased / decreased concurrently while (*V*_*max*_ - *V*_*min*_) remained constant.

**Figure 1.**
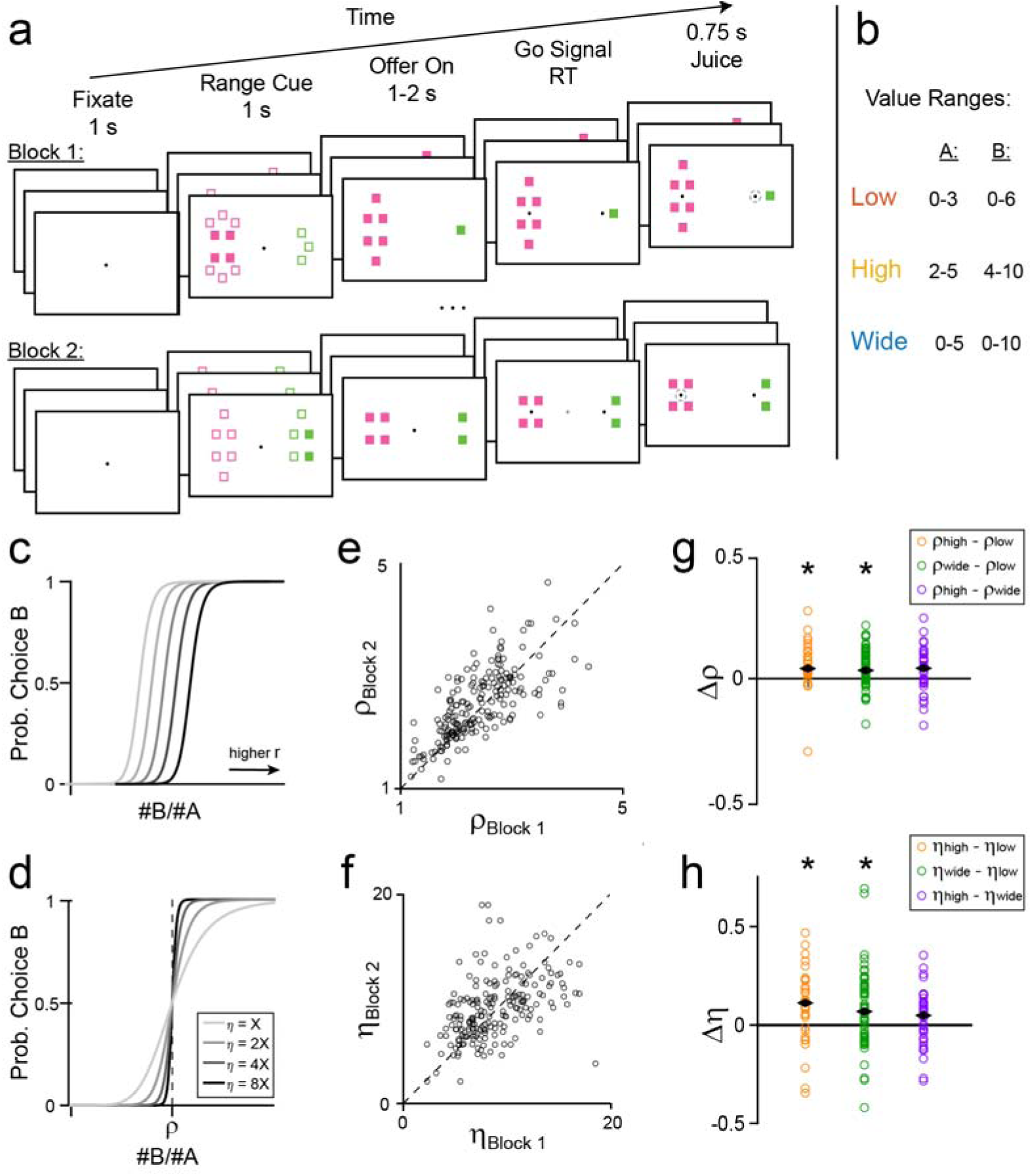
Task outline and behavioral results. **(A)** Schematic of a session. The monkey chose between two juices, each associated with one color. The animal initiates each trial by fixating on a central point. After 1s, range cues appear on either side of the central fixation. Filled squares indicate the minimum possible offer for a given juice and total squares indicate the maximum. In this trial, the cue informed the monkey that the quantity of juice B offered would be between 4 and 10 drops, while the quantity of juice A would be between 0 and 3 drops. After 1s, the cues are replaced by two sets of filled squares representing the current offers. In this trial, the animal chose between 1 drop of grape juice and 4 drops of fruit punch. After a variable interval (1-2s), the central fixation disappears, and targets appear next to each offer, cuing the monkey to indicate its choice. The monkey then makes a saccade to one of the targets and holds fixation for 0.75s, after which it receives the chosen juice. Each session consists of 2-3 blocks, each with ∼250 trials. Ranges remain constant across trials within each block, and change between blocks. **(B)** Each juice is offered in one of three ranges, “low”, “high”, or “wide”. The two juices can be offered in either the same type of range or different types of range within a block. (C-H) Changes in choice behavior across sessions. **(CD)** Illustration of the behavioral response function for changing values of relative value, p (C), or behavioral steepness n (D). Increased p corresponds to a decreased probability of choosing offer B for a given offer (#A:#B). Increased n corresponds to less variable choice behavior. (E-H) Animal behavior across sessions. Each point represents the behavior from one pair of blocks in a session (n=205). **(E)** Relative value in the earlier vs. later block of a session. Values of ρ were strongly correlated across pairs of blocks in a session (r = 0.73, p = 5.5*10-35, Pearson correlation) and slightly elevated in later blocks (median Δp = 0.07; p = 0.01). **(F)** Values of n were correlated across blocks (r = 0.24, p = 4.4*10-4, Pearson correlation) but did not differ between the first and second blocks of a session (p = 0.47, Wilcoxon signed rank test). (G) Fractional difference in p across range types. Median differences: ρhigh-ρlow = 0.040 (p = 2.3*10-4), ρwide-ρlow = 0.032 (p = 8.3*10-5), ρhigh-ρwide = 0.041 (p = 0.54). **(H) F**ractional difference in η across range types. Median differences: ηhigh-ηlow = 0.11 (p = 7.7*10-3), r) η wide-ηlow = 0.069 (p = 4.2*10-3), η high-η η wide = 0.049 (p = 0.14). Black diamonds indicate median **f**ractional differences, and asterisks indicate significant difference from 0 (p < 0.05). All p-values Wilcoxon signed rank test. Fractional difference is defined as the difference in parameter values divided by their sum.

We analyzed the animals’ behavior separately in each trial block. A logistic regression of the choice pattern provided measures for the relative value (*ρ*) and the sigmoid steepness (*η*) (Fig.1D; see Materials and Methods). Choice patterns generally presented a quality-quantity tradeoff between the juices (mean(*ρ*) = 2.4 across sessions). Within a session, *ρ* was strongly correlated across blocks (r = 0.73, p = 5.5*10^−35^, Pearson correlation; Fig.1E), indicating that the juice preferences were fairly consistent within a session. Values of *ρ* increased slightly in the second block compared to the first, presumably reflecting the animals’ increasing satiety: their preference shifted toward the preferred juice rather than the higher quantity (p = 0.01, Wilcoxon signed rank test). Values of *η* were also correlated across blocks (r = 0.24, p = 4.4*10^−4^, Pearson correlation) but did not differ systematically between the first and second blocks of a session (p = 0.47, Wilcoxon signed rank test) (Fig.1F).

Choice behavior was weakly affected by the value range (Fig.1GH). In general, relative values were slightly larger in high and wide range blocks compared to low range blocks (Fig.1G), reflecting an increase in the relative value of A for higher quantities. High and wide range blocks also had steeper sigmoid functions than low range blocks (lower choice variability, Fig.1H). The sigmoid steepness recorded in low range and wide range blocks was statistically indistinguishable (Fig.1H). Differences in sigmoid steepness are likely related to the monkeys’ greater motivation in high value blocks (see Discussion).

### Neural responses adapt to both the maximum and minimum value

We recorded the activity of 1,262 cells from two monkeys as they performed the choice task (monkey D, left hemisphere: 480 cells; monkey F, left hemisphere: 373 cells, right hemisphere: 409 cells). We analyzed the activity of these neurons in seven time windows after offer onset. A “trial type” was defined by two offers and a choice (e.g., [1A:3B, A]). A neuronal response was defined as the activity of one neuron in one time window as a function of the trial type, pooling trial types from two blocks. Building on the results of previous studies (Padoa-Schioppa & Assad, 2006), we identified task-related responses (ANOVA, p < 0.05 in both blocks) and classified them as encoding one of the variables *offer value A*, *offer value B, chosen value*, or *chosen juice* (see Materials and Methods). In total, 488 neurons encoded a decision-related variable in at least one time window (monkey D: 248 cells, 51.7%; monkey F: 240 cells, 30.7%). 1,917 responses passed the ANOVA criterion, and 984 of these encoded the *offer value* or the *chosen value* (Table 1). Of these, 644 value-encoding responses met inclusion criteria for our analysis of neuronal adaptation (see Materials and Methods).

**Table 1.** Number of responses encoding each variable for Monkeys D and F. Laler analyses focused on offer value and chosen value responses. Values in parenthesis indicate the number of responses that met inclusion criteria.

| Response | Monkey D | Monkey F |
| --- | --- | --- |
| *Offer Value A* | 242 (160) | 123 (75) |
| *Offer Value B* | 116 (78) | 88 (51) |
| *Chosen Value* | 249 (173) | 166 (107) |
| *Chosen Juice* | 578 | 355 |

Fig.2 illustrates four potential outcomes for the experiment. First, responses might adapt fully to changes in both maximum and minimum values (range adaptation; Fig.2A). In this case, the slope of encoding would be steeper in the low and high ranges compared to the wide range. In addition, the range of firing rates would be the same in all conditions – the maximum and minimum values in each condition (*V*_*max*_ and *V*_*min*_) would always evoke the same maximum and minimum responses (*R*_*max*_ and *R*_*min*_, respectively). Alternatively, neurons might adapt to the maximum value but not to the minimum value (max adaptation; Fig.2B). Conceptually, this scenario would occur if values were represented relative to the status quo (i.e., the animal’s state prior to the decision). In this case, the encoding slope in the high and wide ranges would be the same, while the slope in the low range would be steeper. In addition, *R*_*min*_ would be elevated in the high value range, reflecting a larger *V*_*min*_. Notably, adaptation to either the value range or the maximum value would be consistent with previous results (Kobayashi et al., 2010; Padoa-Schioppa, 2009). Thirdly, neurons might not adapt at all (Fig.2C). Non-adapting responses would have the same tuning function in all conditions, but different values of *R*_*max*_ and *R*_*min*_ would be observed due to the different values sampled in each range. Since previous work found adaptation to changes in maximum value, we considered this outcome unlikely, but kept it as reference point for our analyses. Finally, neurons might adapt partially to the maximum value, the minimum value, or both (Fig.2D). In this case, value encoding would have a steeper slope for the low and high value ranges relative to the wide range, but the range of evoked responses would also change across conditions. For example, *R*_*max*_ and *R*_*min*_ would be higher in the high range compared to the low range condition, corresponding to higher *V*_*max*_ and *V*_*min*_.

**Figure 2.**
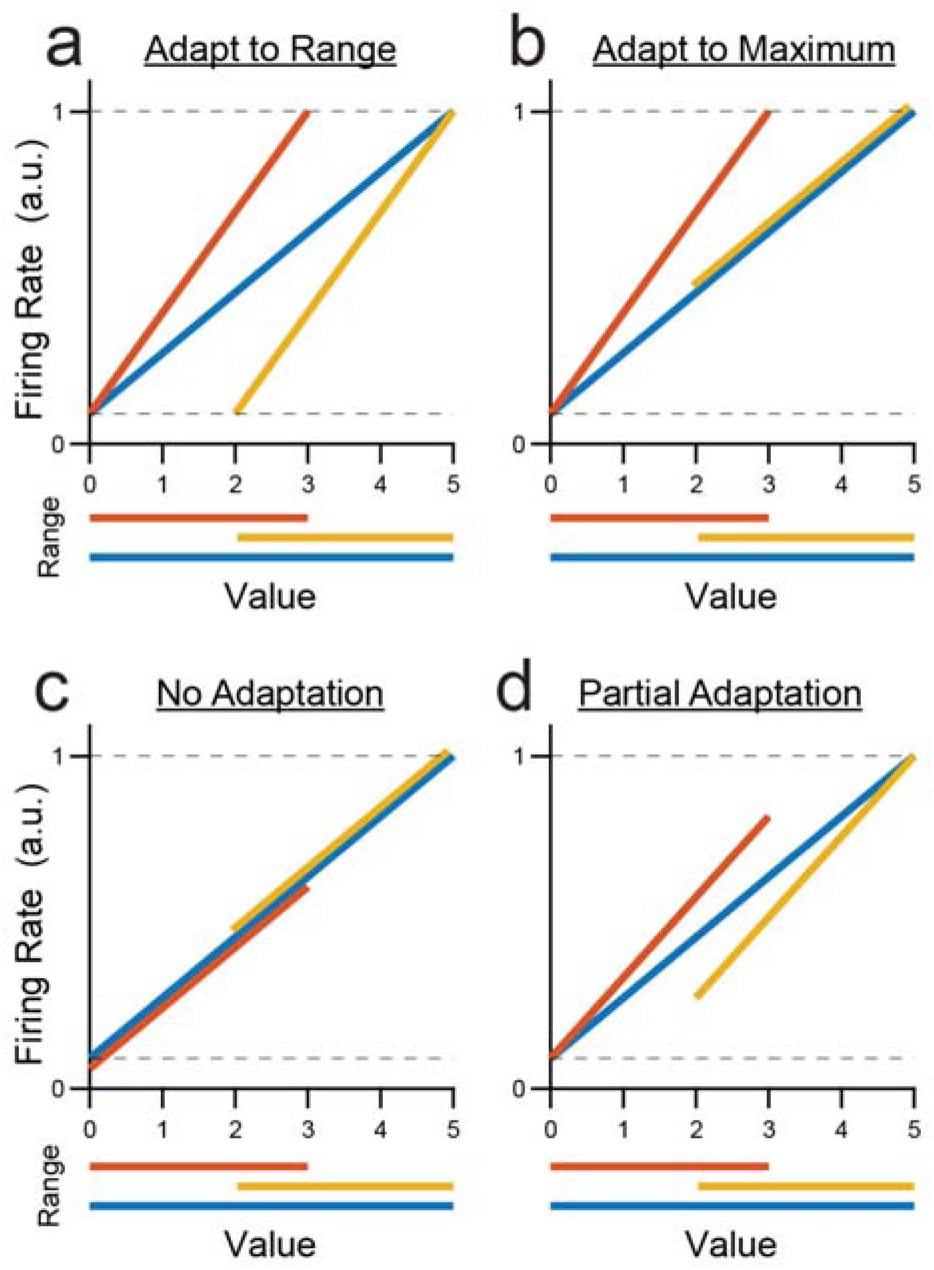
Four hypotheses for neuronal adaptation in a hypothetical offer value A response. Red, yellow and blue traces represent neural responses in the low, high, and wide range conditions. Dotted lines represent the absolute minimum and maximum responses observed for the neuron across all conditions. **(A)** Predicted responses if neurons fully adapt to both maximum and minimum values. The slope of encoding is lower in the wide range (blue) compared to the low range (red) or the high range (yellow). The responses to the minimum and maximum values are consistent across all ranges. **(B)** Predicted responses if neurons adapt to changes in maximum, but not minimum value. Encoding is steeper in the low range compared to the high or the wide range. The maximum response is consistent across conditions, but the minimum observed response is higher when the minimum value is larger (high range). **(C)** Predicted response if neurons do not adapt at all. The response function does not change across conditions. Neurons have the same encoding slope across all ranges. The maximum (minimum) response is higher in blocks where the maximum (minimum) value is larger. **(D)** Predicted responses if neurons partially adapt to changes in both maximum value and minimum value. Encoding is steeper in the low and high ranges compared to the wide, but the full dynamic range is not always used. The maximum observed response is lower when the maximum value is smaller (low range). Similarly, the minimum response is higher when the minimum value is larger (high range)

In broad terms, neurons adapt to a parameter if changing that parameter alters their tuning functions. We frequently observed adaptation in *offer value* and *chosen value* responses for all types of range transition. For example, the cell in Fig.3AB adapted to changes in the maximum value of juice B. It encoded *offer value B* in both blocks, but its tuning slope was shallower when the maximum value increased. Similarly, the cell in Fig.3CD adapted to changes in the minimum value, encoding *offer value A* with a shallower slope when the minimum value decreased. The cell in Fig.3EF adapted to changes in both maximum and minimum value. When the range of chosen values shifted down, the tuning curve shifted left as firing rates rescaled to the new value range. In this case, the encoding slope also increased, reflecting the narrower range of chosen values in the second block.

**Figure 3.**
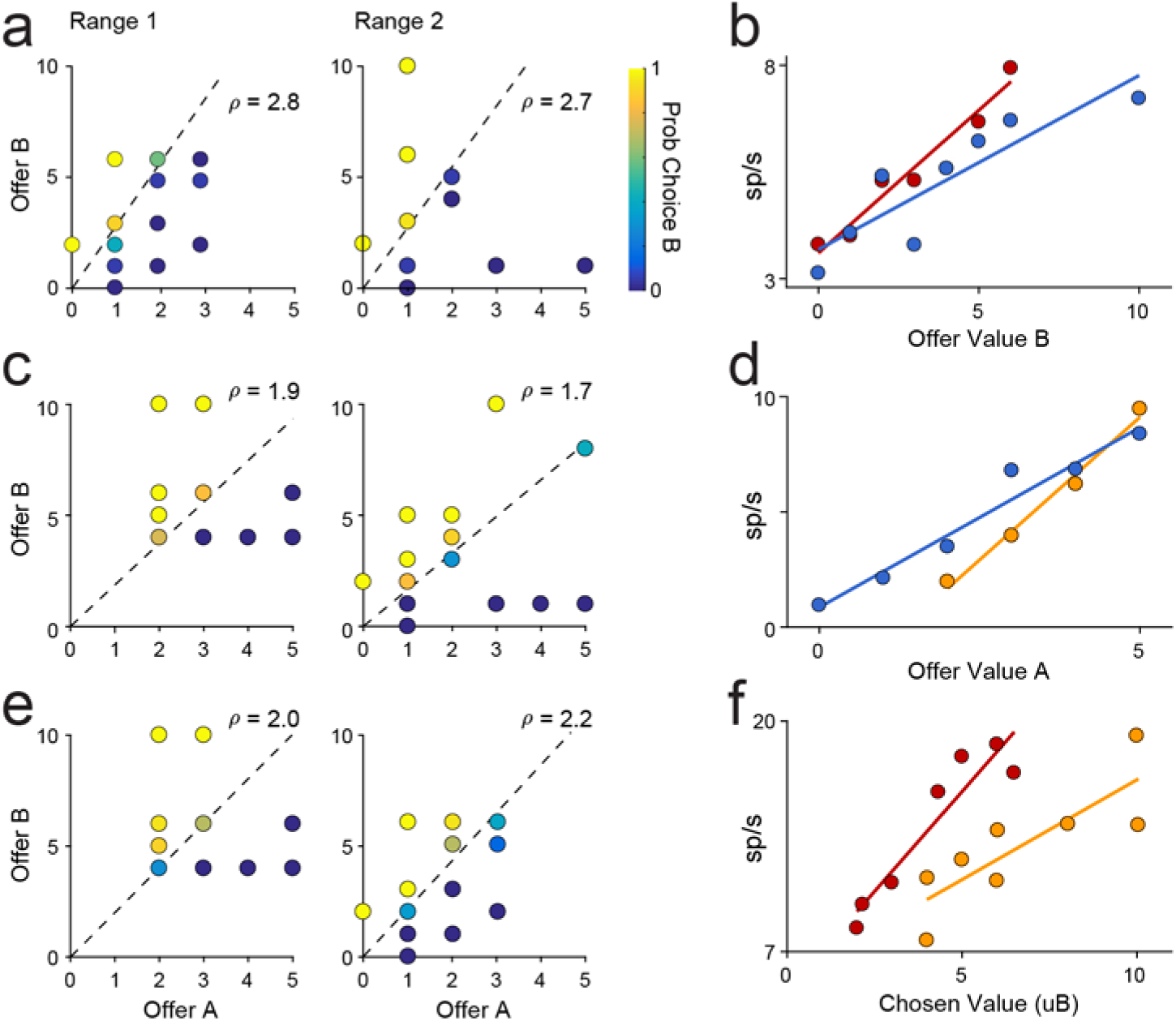
Examples of behavior and neuronal activity for three types of range transition. **(A,B)** Increase in maximum value (low → wide). **(C,D)** Decrease in minimum value (high → wide). **(E,F)** Decrease in both (high → low). (A,C,E) Choice behavior. Each point represents one offer type. Quantity of offer A is shown on the x-axis and quantity of offer B on the y-axis. The color of each point indicates the fraction of trials in which the animal chose juice B. (B,D,E) Neural activity across the two blocks. Colors indicate range type (red = low range; yellow = high range; blue = wide range). Responses encode offer value B (B), offer value A (D), and chosen value.

Across the population, neuronal responses were variable, but they consistently showed adaptation to both the maximum and minimum value (Fig.4A-C). Notably, neuronal adaptation was not complete: the range of firing rates differed across range types, indicating that neural activity did not fully rescale to the range of values available in each trial block. This point can be seen most clearly in Fig.4D. Although each of the three range types have distinct tuning curves, the minimum response is higher in the high range condition compared to the other conditions. Similarly, the maximum response in the low range condition is lower compared to the high and wide range conditions. This result most closely resembles partial range adaptation (Fig.2D).

**Figure 4.**
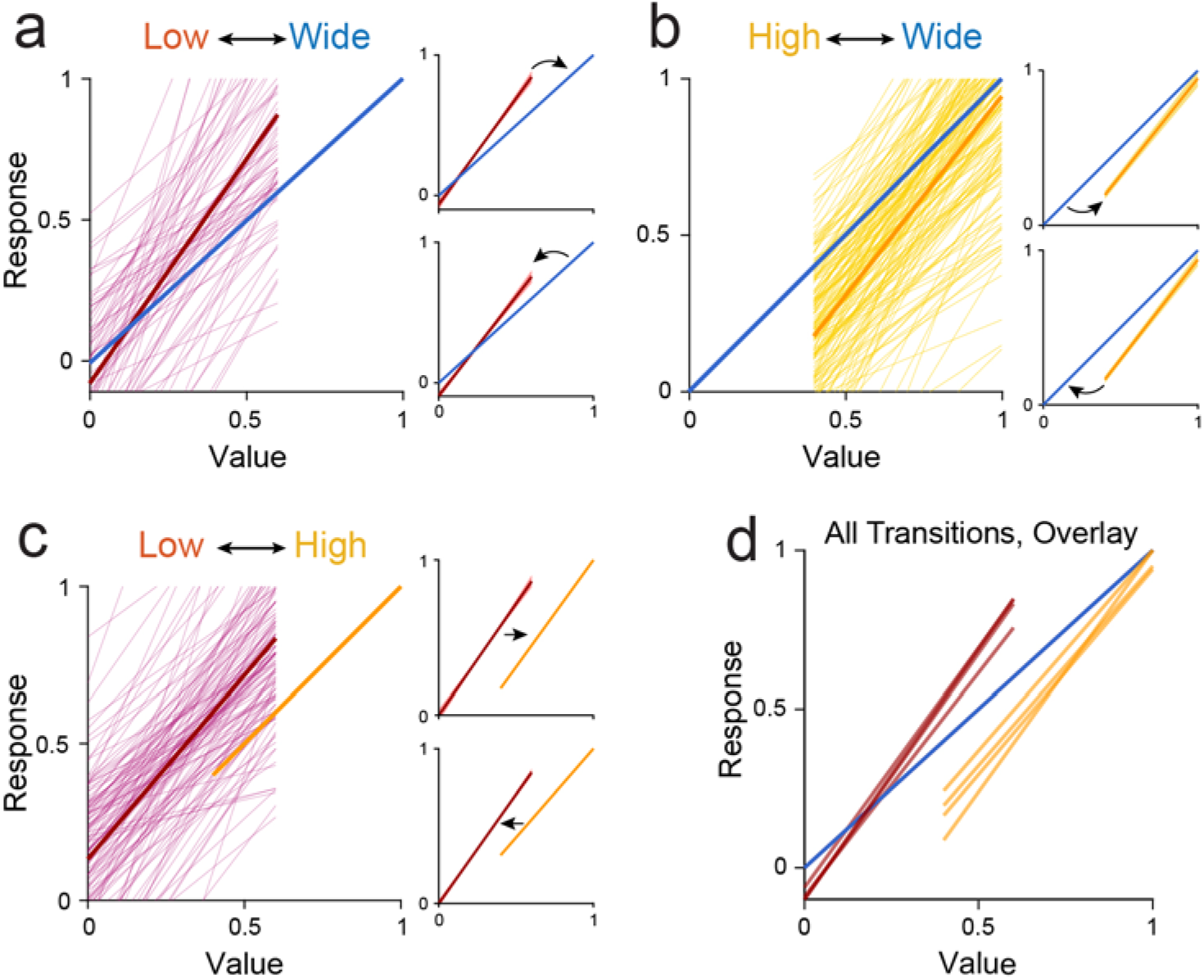
Adaptation in offer value responses across each type of range transition. **(A-C)** Individual responses (thin lines) and population mean (think line) for (A) changs in maximum value (n = 72); (B) change in minimum value (n = 163); and (C) change in both (n = 129). Insets show average responses for transitions where Vmax and/or Vmin increase (top) or decrease (bottom). Shaded region in inset shows mean ± SEM. Responses are normalized to the wide range (A,B) or high range (C). Insets in (C) are normalized to the Vmax(high) - Vmin(low). **(D)** Overlay of mean responses for all six types of range transition. Transitions from (A) and (B) are aligned to wide range. Transitions from (A) and (C) are aligned to the low range.

### Adaptation involves incomplete rescaling

To examine value adaptation quantitatively, we analyzed three features of the response function: the slope of the encoding, the response to *V*_*max*_, and the response to *V*_*min*_.

We analyzed changes in the tuning slope in two ways. First, we compared the slope directly across changes in *V*_*max*_, *V*_*min*_, or both. On average, the slope was larger when the value range was high or low compared to when the range was wide, consistent with the hypothesis that neurons adapt to both maximum and minimum values (Fig.5A-C). Responses also showed slightly higher slopes in the low range relative to the high range condition (Fig.5C). While this observation is consistent with the idea that responses adapt more to *V*_*max*_ than to *V*_*min*_, the effect was driven by *chosen value* responses. *Offer value* responses alone did not show any difference in slope between the low range and the high range conditions. To interpret changes of slope in *chosen value* responses, we also need to account for the difference in value range (*V*_*max*_ - *V*_*min*_), which varies depending on the animal’s choice pattern.

**Figure 5.**
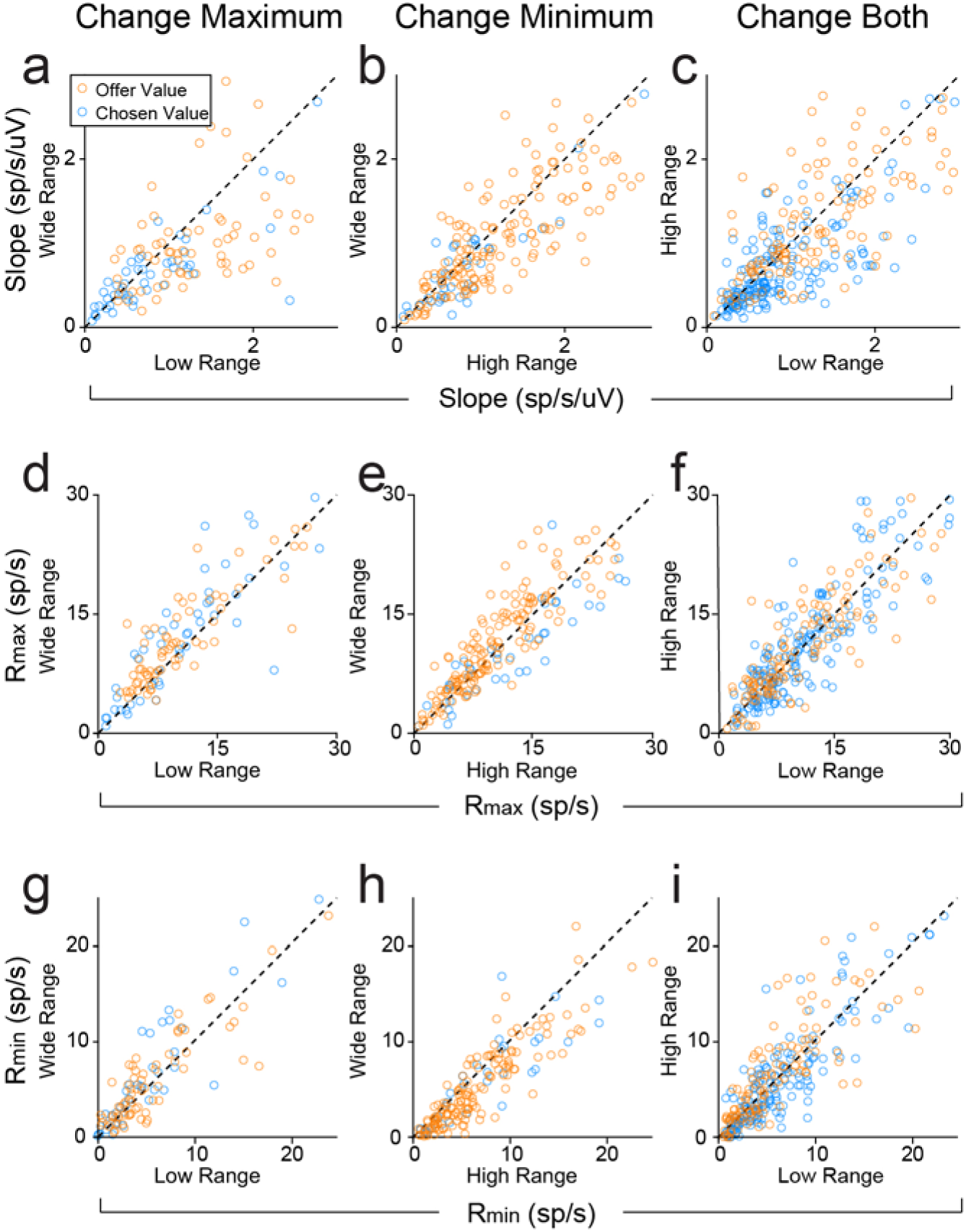
Metrics of value adaptation for each type of block transition. Transition type: maximum value changes (ADG, n = 121), minimum value changes (BEH, n = 206), both change (CFI, n = 317 responses). **(A-C)** Changes in the slope of value encoding of offer value and chosen value responses. Dashed lines show y = x. Value encoding was generally steeper for low or high value ranges compared to the wide range (high vs. wide: p = 1.7*10-7; low vs. wide: p = 7.3*10-8; Wilcoxon signed rank test). When both maximum and minimum values changed, the encoding slope for high and low value ranges was close to the unity line, but slightly higher for the low value range (p = 4.3*10-7, Wilcoxon signed rank test). This effect was driven by chosen value resonses. Offer value responses alone did not show such difference (p = 0.22). **(D-l)** Across-block comparisons of Rmax (D-F) and Rmin (G-l) for each type of range transition. Rmax (Rmin) was generally higher when Vmax (Vmin) was higher. (D) Rmax,wide > Rmax,low (p = 6.7*10-6); 6 points fall outside the limits of the plot. (E) Rmax is not significantly different between high and wide blocks (p = 0.058); 14 points fall outside the limits of the plot. (F) Rmax is higher in the high range compared to the low (p = 5.4*10-4); 16 points outside the limits of the plot. (G) Rmin,wide > Rmin,low (p = 0.026); 6 points fall outside the limits of the plot. (H) Rmin,wide < Rmin.high (p = 6.2*10-9); 9 points fall outside the limits of the plot. (I) Rmin.high > Rmin.iow (p = 9.2*10-4); 8 points fall outside the limits of the plot. Dashed lines show y = x. All p-values based on Wilcoxon signed rank test.

To further examine the relationship between slope and value range, we defined Adaptation Ratios (ARs) for three hypothetical scenarios: adaptation to maximum value (*AR*_*max*_), adaptation to the value range (*AR*_*range*_), or no adaptation (*AR*_*none*_):

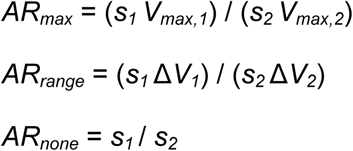

where *s* is the encoding slope, Δ*V* is the value range (*V*_*max*_ - *V*_*min*_), and indices 1 and 2 indicate different trial blocks. For high ⇔ wide or low ⇔ wide transitions, we defined Block 1 as the wide range (ARs are calculated as wide/narrow). For high ⇔ low transitions, we defined Block 1 as the high range (ARs are calculated as high/low). ARs provide a metric for the degree of adaptation. If neurons adapt completely to both maximum and minimum values, then *AR*_*range*_ = 1. If they adapt to the maximum only, then *AR*_*max*_ = 1. Note that *AR*_*none*_ is simply the ratio of slopes in the two conditions, and should be 1 if responses do not adapt. ARs are ambiguous for certain types of range transition, For example, when only the maximum value changes, *AR*_*max*_ and *AR*_*range*_ are equivalent. In addition, ARs only test the relation between the value range and the tuning slope; they are not affected by changes in the intercept of the tuning function. Hence, AR = 1 does not imply that responses adapt in a specific way. However, AR ≠1 indicates that a particular hypothesis *does not* fully describe adaptation.

Table 2 summarizes the ARs for every type of transition. A few results are noteworthy. First, *AR*_*none*_ < 1 for all range transitions, meaning that adaptation occurred consistently. Similarly, *AR*_*max*_ ≠1 for transitions where *V*_*min*_ changed alone or where both *V*_*min*_ and *V*_*max*_ changed, indicating that responses adapted to changes in both maximum and minimum value. At the same time, *AR*_*range*_ > 1 when *V*_*min*_ changed alone and when *V*_*max*_ decreased. This finding indicates that responses did not fully adapt to changes in either *V*_*max*_ or *V*_*min*_. Overall, these results confirm that responses adapted to both the maximum and minimum values, but that the dynamic range did not rescale completely.

**Table 2.** Metrics of adaptation in *offer value* and *chosen value* responses across six types of range transition. Columns 1-3: median Adaptation Ratios calculated for three hypotheses: 1) neurons adapt to max value only, 2) neurons adapt to both max and mill, 3) neurons do not adapt. If a hypothesis is true, AR - 1. Columns 4-5: median normalized difference in R_max_ and R_min_ between blocks. Nonzero values indicate a change in neural activity range between sessions (incomplete adaptation). Asterisks (*) indicate a significant deviation from 1 (columns 1 -3) or front 0 (columns 4-5). Plus (^+^) indicates that the median ΔR_min_ or ΔR_max_ differs from the value predicted for non-adaptive coding. All p < 0.01, Wilcoxon signed rank test.

| *Transition Type* | *AR<sub>max</sub>* | *AR<sub>range</sub>* | *AR<sub>none</sub>* | $\Delta R_{max}$ | $\Delta R_{min}$ |
| --- | --- | --- | --- | --- | --- |
| *Increase Max* | 1.04 | 1.04 | 0.78* | 0.17*,+ | 0.011 |
| *Decrease Max* | 1.13* | 1.13* | 0.74* | 0.22*,+ | 0.060 |
| *Increase Min* | 0.83* | 1.23* | 0.83* | 0.029 | -0.20*,+ |
| *Decrease Min* | 0.88* | 1.39* | 0.88* | 0.018 | -0.19*,+ |
| *Increase Both* | 1.37* | 1.04 | 0.86* | 0.22*,+ | 0.17*,+ |
| *Decrease Both* | 1.27* | 0.94 | 0.82* | 0.17*,+ | 0.19*,+ |

So far, we have examined changes in the gain of value encoding. However, as Fig.4 illustrates, range transitions often led to a shift in the response to *V*_*min*_ (*R*_*min*_) and in the response to *V*_*max*_ (*R*_*max*_). To quantify this effect, we compared *R*_*min*_ and *R*_*max*_ across different ranges (Fig.5D-I). In general, when *V*_*max*_ (*V*_*min*_) was higher, *R*_*max*_ (*R*_*min*_) was also higher (all p < 10^−3^, Wilcoxon signed rank test). Interestingly, *R*_*min*_ was slightly higher in the wide range compared to the low range condition, even though *V*_*min*_ was the same (Fig.5G, p = 0.026, Wilcoxon signed rank test). *R*_*max*_ did not differ significantly between the wide and the high range conditions, although there was a trend toward higher responses in the wide range (Fig.5E, p = 0.058). Importantly, although responses did not remap completely, our results were inconsistent with the hypothesis of no adaptation (Fig.2C). To quantify this point, we computed the normalized change of *R*_*min*_ and *R*_*max*_ (Δ*R*_*min*_ and Δ*R*_*max*_, respectively) and compared them to the values predicted if neurons did not adapt (see Materials and Methods). Δ*R*_*min*_ and Δ*R*_*max*_ were consistently lower than the values predicted for non-adapting cells (Table 2). Along with the analysis of response gain, these results confirm that value-encoding neurons in OFC undergo partial adaptation to changes in the value range.

The observation of partial rescaling in value-encoding responses raised the possibility that adaptation was still ongoing during data collection. An incomplete temporal process could produce the intermediate range adaptation observed in Fig.4. To test this prospect, we computed the tuning function separately in the first and second half of Block 2. If adaptation was temporally incomplete, responses should show greater changes in the second half of Block 2 compared to the first half. Contrary to this prediction, tuning functions for the first and second halves of Block 2 were nearly identical for all transition types (Fig.6). Statistical analyses confirmed that changes in the slope and intercept of the tuning function were present within the first half of Block 2 (all p < 0.01, Wilcoxon signed rank test). Hence, neuronal adaptation occurred relatively quickly after a change in value range, and the features of range adaptation described above reflect the steady state rather than an unfinished transition.

**Figure 6.**
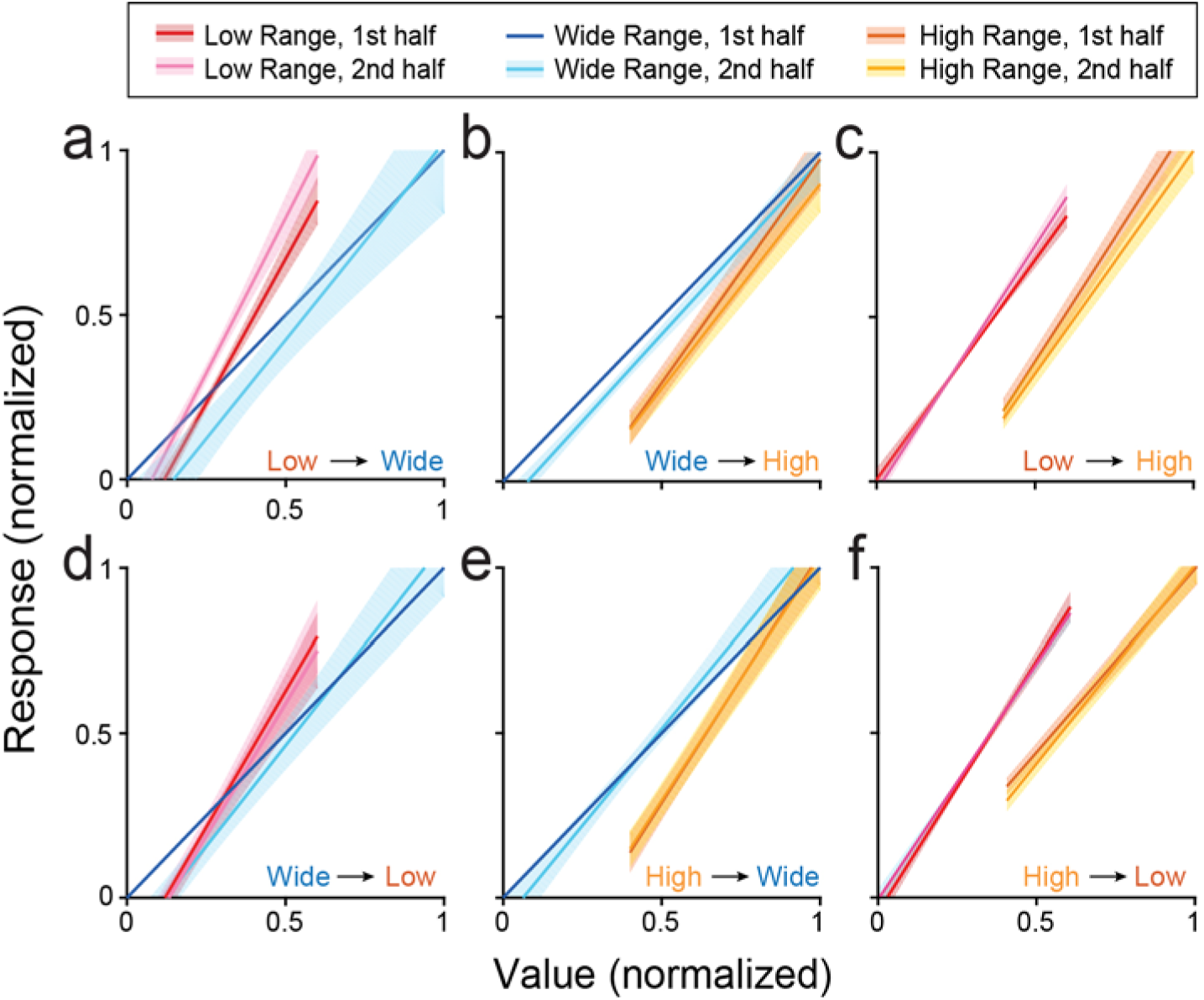
Tuning of offer value responses in the first and second half of each block. Transition types: **(A-C)** Increase in maximum (A; n = 38), minimum (B; n = 64), or both (C; n = 76). **(D-F)** Decrease in maximum (D; n = 33), minimum (E; n = 99), or both (F; n = 53). Shaded regions indicate SEM. The first half of each block is shown in lighter colors, the second half in darker colors. Tuning functions are consistent across the early and late halves of the block.

### Adaptation does not affect linearity of tuning

Previous work found that value encoding in OFC is quasi-linear, but slightly convex on average (Rustichini et al., 2017). We asked whether range adaptation has any effect on this curvature. To address this question, we fit each value-encoding response separately with a quadratic polynomial and a cubic polynomial in each range condition. Confirming previous observations, few responses showed significant quadratic or cubic terms (*β*_2_: 10.6%, *β*_3_: 4.9%; p<0.05, F-test). On average across the population, quadratic terms were slightly positive (p = 5.8*10^−56^, Wilcoxon signed rank test), while cubic terms were slightly negative (p = 1.6*10^−3^, Wilcoxon signed rank test). Most importantly, the distribution of *β*_2_ did not differ between high and low value ranges (Fig.7A; median values: 0.064, 0.61; p = 0.47, Wilcoxon rank sum test). Values of *β*_2_ were slightly lower in the wide range (median: 0.017; p = 9.6*10^−9^ vs. high range, 1.1*10^−10^ vs. low range). However, this difference arose from the fact that the wide range included a greater number of distinct values, which constrained the polynomial fits. Indeed, when we recalculated the quadratic fits for the wide range using only the subset of values present in the low range condition, the distribution of *β*_2_ did not differ from the distribution measured with high and low ranges (median *β*_2,subsampled_ = 0.045; both p > 0.1). Similarly, the distribution of *β*_3_ did not differ across high, low, and wide range conditions (Fig.7B; median values: -0.014, 1.8*10^−3^, and - 3.8*10^−3^; all p > 0.1, Wilcoxon rank sum test).

**Figure 7.**
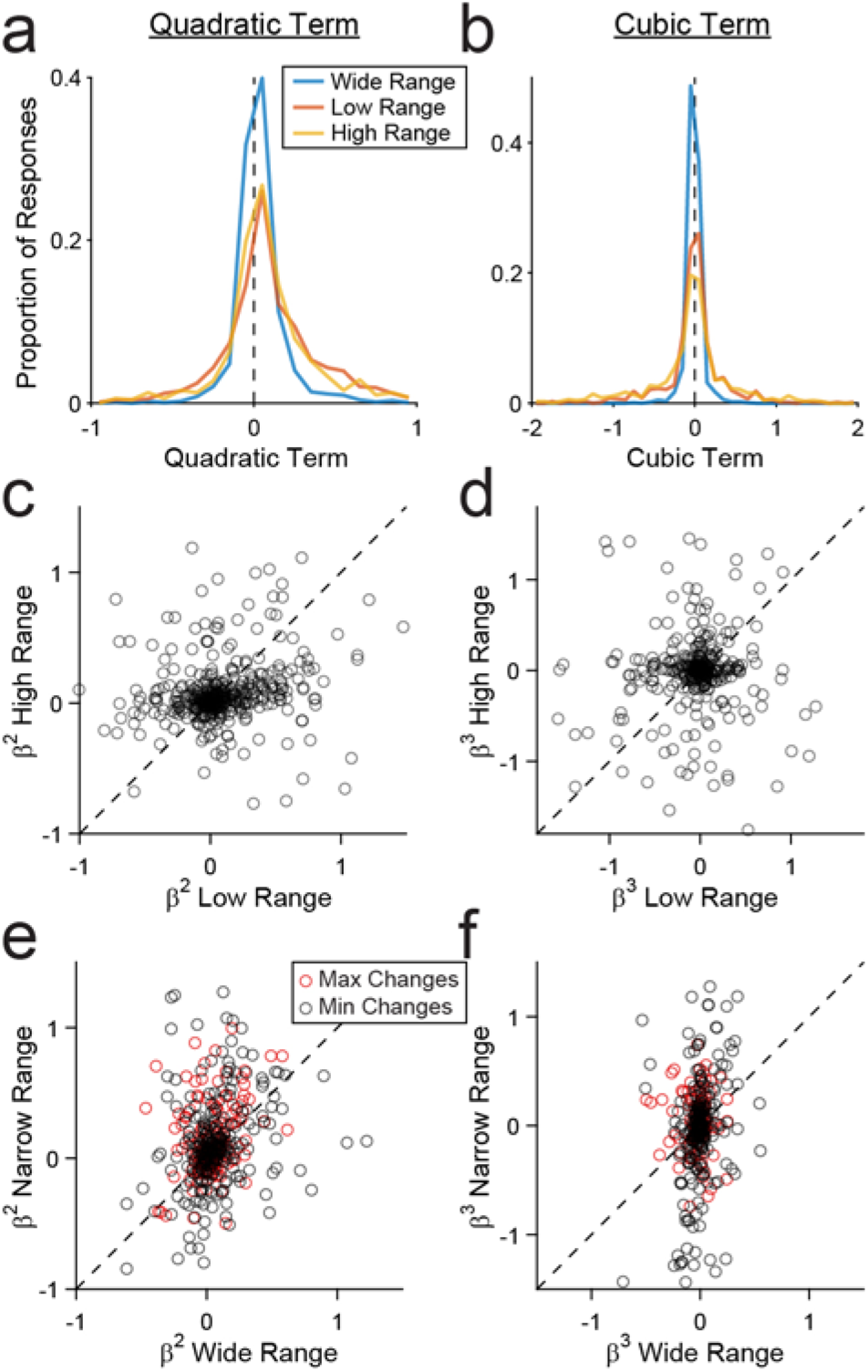
Quadratic and cubic tuning parameters. **(A,B)** Distribution of quadratic (A) and cubic (B) coefficients for all value encoding responses in wide, low, and high ranges. **(C-F)** Quadratic and cubic coefficients for individual responses across blocks. Each point represents one response. Dotted lines show y=x. Correlation (r) and p-values across each transition type: (C) r=0.11, p=1.6 10-3; (D) r = 0.048, p = 0.33; (E) r = 0.39, p = 1.4 10-4 (change in Vmax); r= 0.35, p = 1.1 10-8 (change in Vmin); (F) r = - 0.12, p = 0.12 (change in Vmax); r= 0.24, p = 1.5 10-4 (change in Vmin).

The same pattern of results emerged when we compared *β*_2_ and *β*_3_ for each response across blocks (Fig.7C-F). While values of *β*_2_ varied substantially, coefficients for each response were correlated across blocks. This correlation suggests that *β*_2_ is a characteristic of each neuron’s tuning function. As in the previous analysis, *β*_2_ was slightly higher in narrow ranges compared to the wide range (Fig.7E), although this was only significant for changes in *V*_*max*_ (median difference= 0.031, p = 1.4*10^−4^, Wilcoxon signed rank test). The effect disappeared when *β*_2_ for the wide range was calculated with sub-sampled values (p = 0.39). Values of *β*_3_ did not differ across any type of range transition (all p > 0.1) and did not show any consistent pattern of correlation across blocks.

In summary, adaptation altered the gain and offset of value-encoding responses, but not their quasi-linear functional form.

### Absence of range adaptation in chosen juice cells

All the results presented so far focused on responses encoding the *offer value* or the *chosen value*. In a separate set of analyses, we examined responses encoding the *chosen juice*.

We did not find any evidence of range adaptation in this population. More specifically, we did not find systematic differences in the encoding slopes (difference in responses to preferred and non-preferred juice) or in the minimum responses, across any range transition (Fig.8, all p > 0.05, Wilcoxon signed rank test). Thus it appears that *chosen juice* responses, capturing the binary choice outcome, are not affected by changes in the value range.

**Figure 8.**
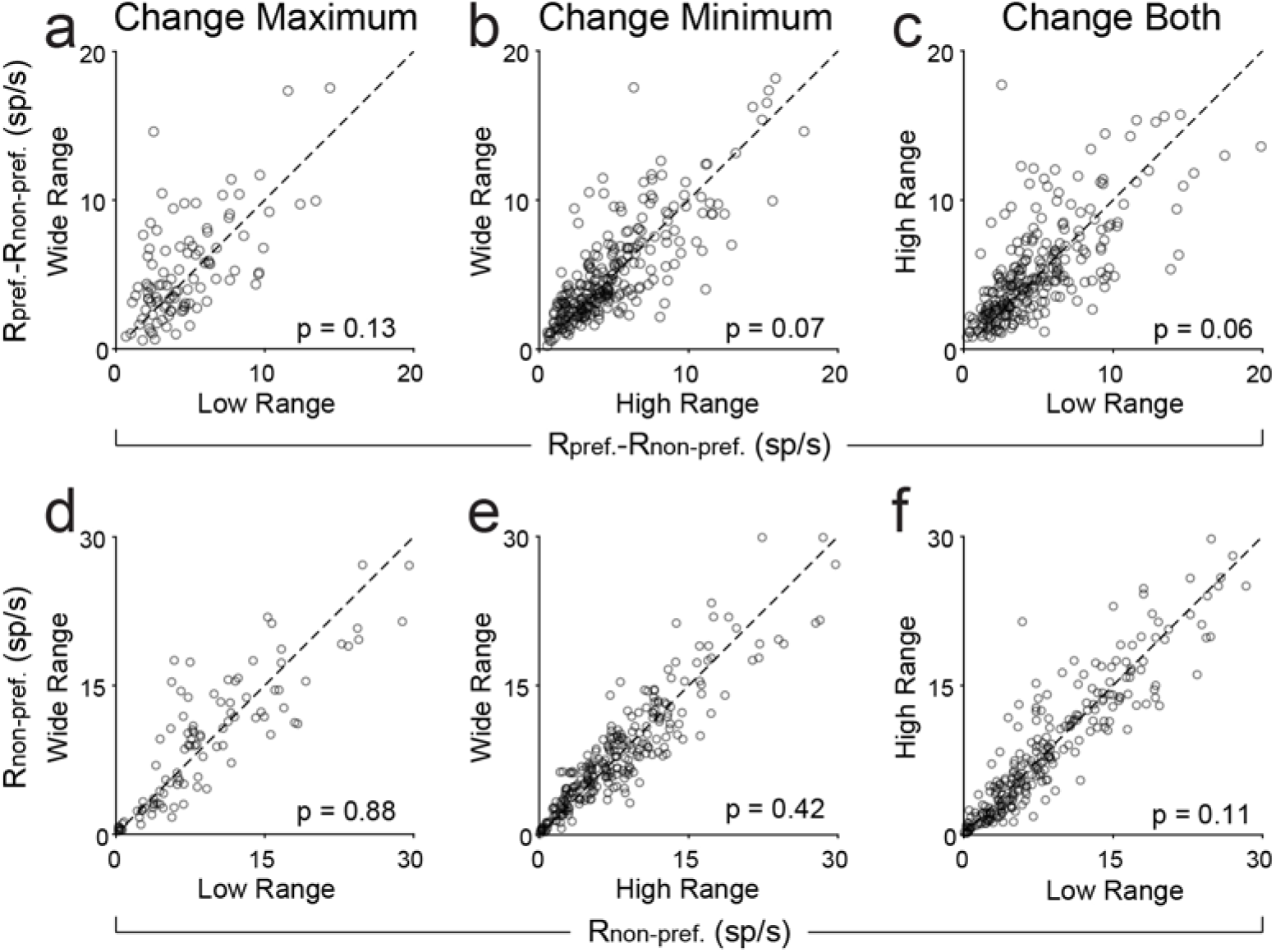
Chosen juice responses to not adapt changes in range. **(A-C)** Slope of chosen juice encoding. **(D-F)** Y-intercept (response to the non-preferred juice) for chosen juice responses. Ranges were defined as the range of the preferred juice (i.e. ranges of juice A were used fora chosen juice cell that fired more for choice A). Defining ranges by chosen value or total value range did not alter results. Dashed lines show y = x. All p-values based on Wilcoxon signed rank test.

### Non-zero baseline activity in offer value cells impairs simulated choice behavior

We have shown that value-encoding neurons do not rescale completely to changes in value range. In other words, responses do not span the full range of potential firing rates in every condition. One important question is whether and how partial adaptation in *offer value* cells affects economic decisions. This issue is closely related to that of optimality in the neuronal representation of subjective values.

In sensory systems, “optimal tuning” generally refers to the neuronal response function transmitting maximal information about the stimuli (Barlow, 1961; Laughlin, 1981). In the neural system underlying economic decisions, this concept of optimality seems less relevant. Instead, optimal tuning may be defined as the response function that maximizes the expected payoff (Rustichini et al., 2017). In our choice task, the payoff is simply the value chosen by the monkey on any given trial. Notably, while the relative value of two juices is subjective, the payoff of two options may be compared objectively once the relative value of the juices is known. For example, if the choice pattern indicates that *ρ* = 2.6, then the payoff of 3B is higher than the payoff of 1A. Importantly, the expected payoff is inversely related to choice variability. When choice variability is higher – i.e. when decisions between two options are more frequently split – the animal is more likely to choose the lower value (lower expected payoff). In previous computational work, we found that a decision network achieved the maximum expected payoff if *offer value* cells adapted completely to the value range – in other words, if their dynamic range rescaled fully to the current range of values (Rustichini et al., 2017). However, that study only considered changes in the slope of the encoding. Moreover, the analysis was limited to instances where the minimum offer value was zero, and it assumed that the response to the minimum offer (i.e., the baseline activity) was also zero. Contrary to these assumptions, here we found that value-encoding responses adapt to the minimum as well as the maximum value. Furthermore, their baseline activity is non-zero and varies systematically with the value range.

To explore the behavioral implications of non-zero, context-dependent baseline activity, we ran a series of computer simulations. We examined a linear decision model comprised of 5,000 *offer value A* and 5,000 *offer value B* units (see Materials and Methods). Each unit encoded the value of its preferred juice in a linear way. Trial-to-trial variability was correlated across units, with correlation values estimated based on empirical measures (Conen & Padoa-Schioppa, 2015). We simulated the choices of this network between pairs of offer values, which were randomly selected on each trial. The decision was determined based on the activity of the two pools of *offer value* cells. Thus, on trials where the activity of *offer A* units exceeded that of *offer B* units, juice A was chosen (and vice versa).

We examined the choice pattern of this network as the minimum activity level (*R*_*min*_) varied. We specifically considered two scenarios. (1) Each unit had a fixed *R*_*max*_, such that increasing *R*_*min*_ reduced the available dynamic range (Fig.9A). (2) Each unit had a fixed activity range (Δ*R = R*_*max*_ – *R*_*min*_), such that increasing *R*_*min*_ shifted the dynamic range (Fig.9B). In essence, the first scenario captures the case where neurons do not adapt to changes in the minimum value; the second scenario is analogous to the partial range adaptation observed in the experiments, where both *R*_*min*_ and *R*_*max*_ are elevated when the value range shifts up (e.g. Fig.4C). For each scenario, we simulated choices for increasing levels of *R*_*min*_. Furthermore, we quantified the effectiveness of choice behavior using the fractional lost value (FLV), defined as:

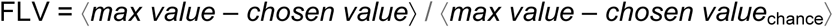

where *max value* refers to the higher value of the two offers on a given trial, *chosen value*_chance_ is the average of the two offers, and ⟨ ⟩ indicates an average across trials. Notably, if a subject always chooses the *max value*, FLV = 0; if the subject always chooses randomly, FLV = 1.

**Figure 9.**
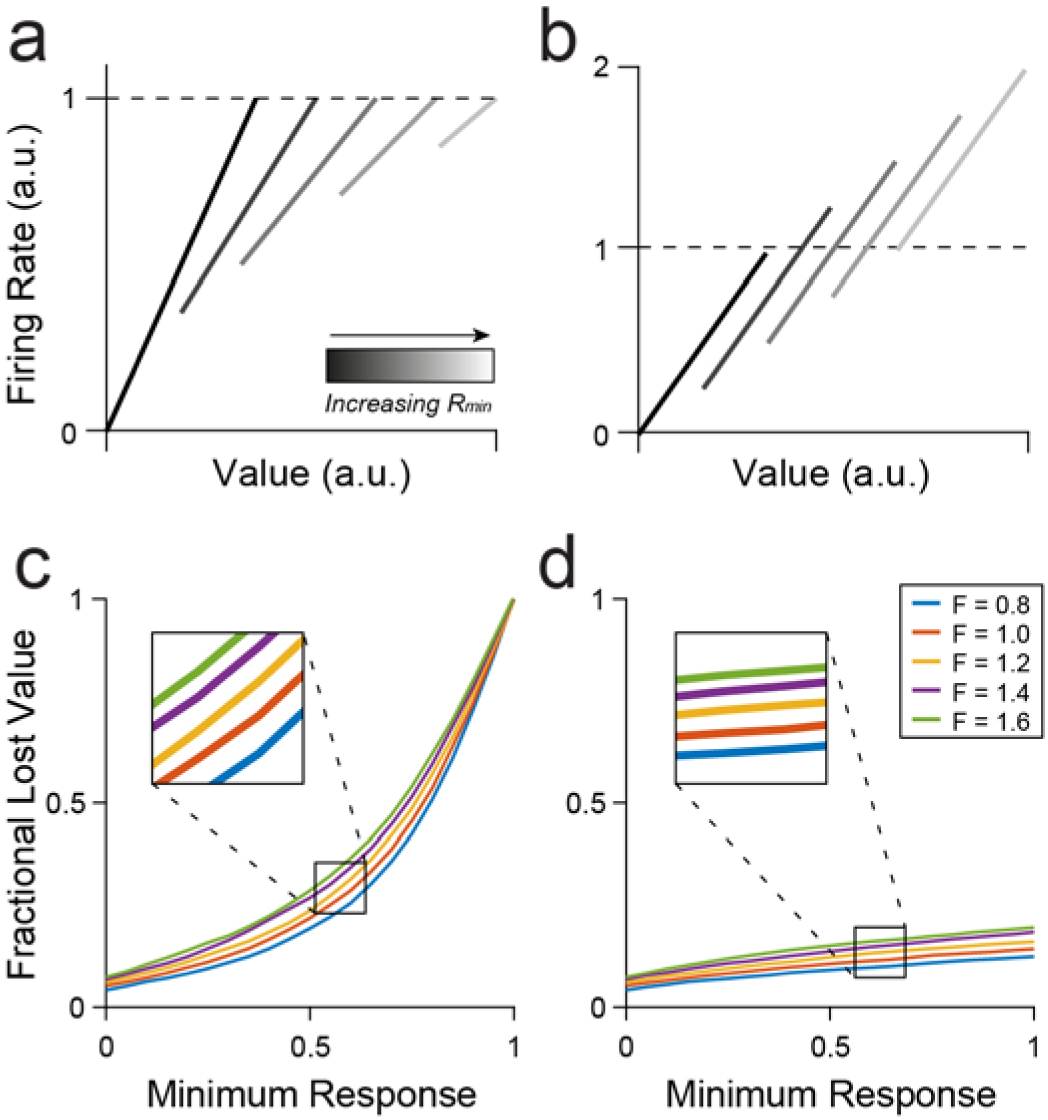
Choice simulation. Fractional lost value (FLV) increases with increasing Rmin. **(AB)** Illustration of example response functions. (A) Rmin increases while Rmax remains fixed. Analogous to a scenario where units adapt only to the maximum value and the value range shifts higher. (B) Rmin increases while the value range (Rmin - Rmin) remains fixed. Analogous to the activity offset associated with partial range adaptation in OFC responses. **(CD)** Simulation results. (C) For neurons with a fixed maximum value, FLV increases to 1 (chance level) as the baseline response increases. (D) For neurons with a fixed range, FLV increases mildly as baseline activity increases. Trace colors indicate results simulated for different Fano factors. Each curve covers 100 values of Rmin. and shows the mean of 20 simulated sessions for each value of Rmin.

Fig.9CD illustrates our results. The payoff decreased with increasing values of R_min_ in both scenarios. However, the presence of a baseline firing rate was much more costly when *R*_*max*_ was fixed (Fig.9C). In the first scenario, FLV increased to 1 as *R*_*min*_ → *R*_*max*_, reflecting the gradual loss of dynamic range. In contrast, the increasing baseline had a much milder effect when *R*_*max*_ and *R*_*min*_ increased together (Fig.9D). In this condition FLV < 0.25 even for *R*_*min*_ equal to or exceeding the total response range.

In summary, increasing the baseline response moderately decreases the expected payoff. However, reducing the dynamic range has a far greater cost.

## Discussion

We showed that value-encoding neurons in OFC adapt to changes in both the maximum and the minimum value available in any behavioral context. Notably, while responses showed consistently higher gain in blocks with a narrow (high or low) value range, neural activity range did not rescale completely to the current value distribution. The range of firing rates was lower when the range of values was lower. Thus value encoding fell in an intermediate zone between fully adaptive coding (range adaptation) and absolute value coding (no adaptation). Importantly, this result did not reflect an unfinished process of adaptation, as tuning functions reached a steady state within the first half of each trial block.

Our results resonate with previous observations. Kobayashi et al. (2010) measured range-dependent changes in value-encoding neurons in several sub-regions of OFC. Their analysis focused on changes in gain. While they divided neurons into adapting, non-adapting, or partially adapting groups, their results are also consistent with a single population of partially adapting responses. Along similar lines, in human subjects, Burke et al. (2016) found partial adaptation in the BOLD signal in ventromedial prefrontal cortex (vmPFC) using a decoding approach. Taken together, these findings suggest that partial adaptation may be a common characteristic of value coding in prefrontal cortex.

The present study resolves an important ambiguity in our understanding of value coding. We showed that OFC neurons adapt to the value range rather than to the maximum value alone. In other words, values are not encoded relative to the subject’s pre-decision state. Instead, values are represented in terms of the best and worst possible outcomes in the current behavioral context. In addition to this insight, our work highlights the importance of analyzing baseline neuronal responses, which are often ignored for the sake of simplicity. Indeed, we have shown that the baseline activity in OFC changes systematically in ways that may affect choice behavior.

### Offsets in the activity range are inefficient

In a previous study, a simulated decision network yielded the highest payoff when neurons exploited their full dynamic range (Rustichini et al., 2017). Here, we found that responses do not span their entire dynamic range in all conditions. Moreover, response functions shift up or down depending on the value range, which we describe as a change in offset or baseline activity. In a simulated decision network, higher baseline activity reduces the expected payoff. While this effect was strongest when the baseline restricted the dynamic range, higher baseline responses increased FLV even when the maximum response also increased. Intuitively, this inefficiency arises from the fact that the variance of neural responses scales with the mean. Ceteris paribus, when a neuron’s dynamic range is higher, firing rates are noisier.

Given the potential cost of a larger response offset in high value ranges, it is worthwhile to speculate on the origins and possible benefits of this phenomenon. One possibility is that neurons adapt to the range of received values rather than to the range of offer values. This interpretation is supported by results from an fMRI study that found that the BOLD signal in vmPFC adapted to the range of received – but not observed – outcomes (Burke et al., 2016). However, this interpretation only accounts for partial adaptation to the minimum value. It cannot explain the change in response to the maximum value or the fact that intermediate adaptation was also found in *chosen value* responses.

Another possibility is that value adaptation may be affected by the overall task structure. In our experiments, monkeys were highly trained on the range adaptation task, and they were familiar with all possible transitions between high, low, and wide ranges. While complete adaptation would warrant an efficient representation of values within a block, it would also limit the circuit’s ability to respond when the value range changes. In contrast, intermediate adaptation reserves a portion of the dynamic range for new values that may appear after a transition. This interpretation suggests that value encoding depends on at least two components: a slow, learning-based process that draws on contextual knowledge; and a more rapid adaptive component that adjusts to the locally experienced value range.

Finally, intermediate adaptation may allow the circuit to maintain information about the overall value of the current context (i.e. the value of the block). Information about the current contextual value makes it possible to predict future reward expectations and affects subjects’ motivation to engage in the task. Moreover, effective value comparison in an adapting network requires information about the distribution of available values as well as neural activity levels on a given trial. Without some mechanism for maintaining this information, signals are ambiguous across contexts and cannot guide behavior effectively (Fairhall, Lewen, Bialek, & De Ruyter van Steveninck, 2001; Rustichini et al., 2017). The differences in response offset observed in OFC may be used by the network to help distinguish the current value state.

### Possible mechanisms of value adaptation

Although our study did not investigate the physiological mechanism of adaptation directly, a few possibilities may be considered. We showed that value adaptation involves both an additive and a multiplicative component. While adaptation to the maximum can occur via a simple change in gain, adaptation to the minimum requires both a change in gain and a horizontal shift in the response function. When the difference between maximum and minimum values is constant, adaptation is purely horizontal: the slope of neuronal encoding remains the same, but responses remap to a new set of values. Additive changes in activity often arise from changes in hyper-polarization or shunting inhibition (Chance, Abbott, & Reyes, 2002; Holt & Koch, 1997). Alternate explanations, such as cell-intrinsic changes in membrane conductivity, generally involve a mixture of additive and multiplicative effects, which is difficult to reconcile with the purely additive adaptation we observed during high-to-low range transitions (M V Sanchez-Vives, Nowak, & McCormick, 2000; Maria V Sanchez-Vives, Nowak, & McCormick, 2000). The multiplicative component of value adaptation could arise from several potential mechanisms. Changes in gain can be produced by both cell-intrinsic mechanisms, such as changes in ionic conductance (Díaz-Quesada & Maravall, 2008; Higgs, 2006; Mease, Famulare, Gjorgjieva, Moody, & Fairhall, 2013), and by circuit-level changes in inhibitory activity (Natan, Rao, & Geffen, 2017; Olsen, Bortone, Adesnik, & Scanziani, 2012; Wilson, Runyan, Wang, & Sur, 2012) or the background level of synaptic activity (Chance et al., 2002). Short-term depression (STD) can also induce changes in gain. Although STD generally has a time constant of a few hundred milliseconds, a longer component lasting tens of seconds has also been observed (Kohn, 2007; Varela et al., 1997).

Recent work examining a more medial region of OFC found that adaptation to simultaneously presented values was best explained by a divisive normalization model (Yamada et al., 2018). The data from our study, which reflect a slower form of adaptation across trials, do not appear to follow a similar model. Among other features, the divisive normalization model predicts a decrease in the maximum response in conditions with a higher value range, which we do not observe. Notably, that experiment focused on adaptation on a very short time scale (∼ 100 ms). Another recent model combined slow and fast normalization dynamics to explain variability in choice behavior across contexts (Zimmerman, Glimcher, & Louie, 2018). One interesting question is whether this model can also account for the neuronal responses recorded in OFC. Divisive normalization is a common form of adaptation in sensory regions (Beck, Latham, & Pouget, 2011; Ohshiro, Angelaki, & DeAngelis, 2011; Olsen, Bhandawat, & Wilson, 2010; Valerio & Navarro, 2003; Wark, Lundstrom, & Fairhall, 2007), and it is highly effective at maximizing the transmission of sensory information across a wide variety of stimuli (Carandini & Heeger, 2011; Simoncelli & Schwartz, 2001). At the same time, divisive normalization seems less well suited for contextual adaptation in a decision circuit, which ideally would optimize the choice outcome rather than transmitting maximal information about the value distribution (Rustichini et al., 2017). Nevertheless, the possible reconciliation of divisive normalization and range adaptation remains an open question.

### Discrepancies in behavioral results

Our behavioral analyses revealed range-dependent changes in both the relative value and the sigmoid steepness (Fig.1). The increased relative value in high-value blocks could be explained if the value of additional juice decreases at higher quantities (diminishing marginal utility). Since A is generally offered in lower quantity, such a nonlinearity would presumably shift preferences toward A when the offer quantities increased. The changes in steepness were somewhat more surprising. A recent analysis of behavior across different ranges found that decision patterns were generally noisier during blocks with higher maximum values, consistent with the idea neurons that encoded value with lower resolution during these blocks (Rustichini et al., 2017). In addition, in our simulations, units with higher baseline activity (analogous to the high value range) produced noisier choice behavior. Yet, in the experiments, the sigmoid steepness changed in the opposite direction (steeper choice functions with the wide and high value ranges compared to the low range). The reason for this discrepancy is unclear, but it may partially reflect the monkeys’ greater motivation during more rewarding blocks. Consistent with this idea, choices were least variable in the high-value range, slightly more variable in the wide range, and most variable in the low range. To shed more light on this issue, future work should carefully match the reward rate across blocks.

To conclude, we examined how the neuronal representation in OFC adapted to changes in maximum and minimum of the value distribution. We found that both maximum and minimum values influence the gain of value encoding, but only partially, leading to an offset in neuronal activity levels across ranges. Theoretical considerations suggest that partial (as opposed to full) adaptation should negatively affect choices. Future work should test this prediction.

## Materials and Methods

All experimental procedures conformed to the NIH Guide for the Care and Use of Laboratory Animals and were approved by the Animal Studies Committee at Washington University in St. Louis. Two adult male rhesus macaques (*Macaca mulatta;* D, 11.5 kg; F, 11.0 kg) were used in the study. Before training, a head-restraint device and a recording chamber were implanted on the skull under general anesthesia. The recording chamber (main axes, 50 x 30 mm) was centered on inter-aural coordinates (A30, L0). Structural MRI scans were obtained before and after implantation and used to guide recording.

### Range adaptation task

In this experiment, monkeys performed a variant of a juice choice task used in several previous studies (Padoa-Schioppa & Assad, 2006). The task was run on custom-written software (http://www.monkeylogic.net/) based on Matlab (MathWorks). Eye position was monitored with an infrared video camera (Eyelink; SR Research). During the experiments, the monkey sat in an electrically insulated enclosure (Crist Instruments) with its head fixed. Cues were displayed on a computer monitor placed 57 cm in front of the animal.

Monkeys chose between two juices, A and B, offered in varying quantities. Juice A was defined as the preferred juice (i.e. 1A was generally chosen over 1B). On each trial, the monkey began by fixating on a central point. After 1s, cues appeared on each side of the central fixation, indicating the current range of possible offers. The cues consisted of a set of filled and empty colored squares. The color of the squares indicated the juice type, the total number of squares represented the maximum possible offer for that juice in the current trial, and the filled squares represented the minimum possible offer in that trial (Fig.1A). The cues remained on screen for 1s and were then replaced by a set of solid squares denoting the offers on the current trial. After a randomly variable delay (1-2 s), the central fixation point disappeared and targets appeared next to each offer (go signal). The monkey indicated its choice with a saccade to one of the targets and, after 0.75 s, received the juice corresponding to the chosen offer. If the monkey broke fixation before the go signal appeared or if he failed to fixate the target for 0.75 s after the saccade, the trial was aborted and the monkey received no reward.

Each session consisted of 2-3 blocks, each lasting ∼ 250 trials. The offered quantity varied pseudo-randomly from trial to trial within a defined range. Within a block, the range of possible offers was kept consistent for each juice. The monkey could either learn the value range implicitly through experience or explicitly by use of the range cues. We do not attempt to distinguish between these possibilities here. Between blocks, the range of available offers for each juice changed, with three possible ranges for each juice: “high” (2-5 units of juice A or 4-10 units of B), “low” (0-3 uA or 0-6 uB), and “wide” range (0-5 uA or 0-10 uB). Most range transitions consisted of an increase/decrease in the minimum value (*V*_*min*_) while the maximum value (*V*_*max*_) either remained constant or shifted in conjunction with *V*_*min*_. Note that when *V*_*min*_ and *V*_*max*_ changed together, the difference *V*_*max*_ - *V*_*min*_ was kept constant. We counterbalanced the type of range transition across sessions. In a smaller subset of sessions, *V*_*max*_ increased/decreased while *V*_*min*_ was kept at zero. The ranges of juice A and B could change in either the same direction or different directions in a given session.

### Analysis of behavior

All analyses were conducted in Matlab (MathWorks). Unless otherwise noted, reported p-values were calculated using the Wilcoxon signed rank test. Choice behavior was analyzed separately for each block. We defined the choice pattern as the percent of trials in which the animal chose juice B as a function of the offer ratio (#B/#A). We fit the choice pattern to a sigmoid function using logistic regression:

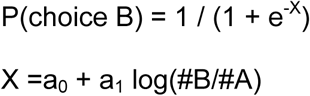

From this fit, we computed the relative value of the two juices (*ρ*) and the sigmoid steepness (*η*):

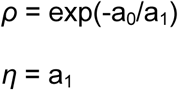

We examined changes in *ρ* and *η* as a function of range type. To do so, we compared data for all pairs of blocks within a session. We recorded during 107 sessions, each of which included 2-3 range conditions, yielding a total of 236 unique block pairs. Block pairs were excluded from the behavioral analysis if there were <2 offer types with choices split between the two juices (31 block pairs excluded). If there are <2 split offer types, a range of parameters can fit the data equally well, making it impossible to precisely identify *ρ* and *η*. For the remaining 205 block pairs, we computed a fractional difference for each parameter across different range types, where we defined fractional difference the value difference divided by the value sum.

### Electrophysiology

We recorded neuronal data from the central OFC of two monkeys, in a region approximately corresponding to area 13m (Ongur & Price, 2000) (monkey D: A 31:36, L -6:-10; monkey F: A 31:37 L -6:-11 and 6:11). Recordings were obtained using tungsten electrodes (125 µm diameter; FHC) and 16-channel silicon V-probes (185 µm diameter, 100 µm spacing between electrodes; Plexon). Electrodes were lowered vertically into position each day using a custom-built micro-drive (step size: 2.5 µm). Recording depth was determined ahead of time based on structural MRI.

Electrical signals were amplified (gain: 10,000) and band-pass filtered (low-pass cut-off: 300 Hz, high-pass cut-off: 6 kHz; Lynx 8, Neuralynx). Action potentials were detected on-line by setting a threshold during recording, and waveforms crossing the threshold were saved (40 kHz sampling rate; Power 1401, Cambridge Electronic Design). Spike sorting was conducted off-line using standard software (Spike 2, Cambridge Electronic Design). Neurons were included in the analysis if they remained stable and well-isolated in two blocks for at least 120 trials per block. Responses that were not stably isolated for the full session were only analyzed for the trials in which they were stable. In the V-probe recordings, spikes from the same neuron were occasionally picked up by two neighboring contacts. These were detected manually based on the consistent presence of simultaneous spikes. If units in neighboring channels shared >70% of spikes, they were considered the duplicates and one of the units was excluded from analysis.

### Response classification

We analyzed cell data in seven time windows following offer onset: post-offer (0.5 s after offer onset), late-delay (0.5-1.0 s after offer onset), pre-go (0.5 s before the go signal), reaction time (time from go cue to target acquisition, usually ∼ 200ms), post-juice (0.5 s after juice delivery) and post-juice 2 (0.5 s to 1s after juice delivery). Data were analyzed independently for each block. We defined a “trial type” as a set of two offers and the monkey’s choice between them. For example, if the monkey chose B on a trial where he was offered 1A vs. 6B, the trial type would be [1A : 6B; B]. Task-based neuronal activity was calculated by taking the mean firing rate for each trial type in each time window. A “neuronal response” was defined as the activity of one cell in one time window across two blocks. Since we were interested in the effects of adaptation at steady state, we discarded the first 16 trials of each block before analysis, thereby excluding trials where the monkey had not yet experienced the full range of values.

A response was considered task-related if it passed an ANOVA (factor: trial type; p < 0.05) in both blocks. To classify task-related responses, we regressed each response against the variables *offer value A, offer value B, chosen value,* and *chosen juice*. Regressions were performed separately for each of the two blocks. We classified a response as encoding a variable if 1) the regression on that variable had a nonzero slope in both blocks (p < 0.05) and 2) in cases where more than one option met the first criterion, that variable had the highest total R^2^ in the two blocks. Further analyses focused on *offer value* and *chosen value* responses. Since we were interested in the effects of changing the value distribution, we excluded responses from analysis if the value range differed by <0.5 units of value between blocks (132 responses). We also excluded cells with dramatic changes in pre-trial firing rate (>1.6x change during the fixation time window, 208 responses), since large variability in baseline activity could obscure effects on cell tuning. Including these responses in the analysis added noise but did not qualitatively alter the results.

Most neuronal responses encoded value with a positive slope (i.e. firing rates increased with value, 71% of responses). For our analyses, we rectified negative encoding responses and pooled all responses. The goal of rectifying responses is to maintain the same range of responses and same slope magnitude, but with a positive rather than negative sign. We rectified the slope (*s*) and intercept (*b*) of negative encoding responses as follows:

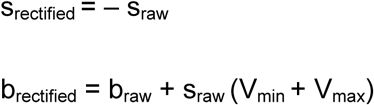

With this approach, the rectified response covers the same range of firing rates as the original, but the maximum evoked response now corresponds to *V*_*max*_ rather than *V*_*min*_. We confirmed that analyses produced qualitatively similar results for positive and negative encoding responses. Restricting the analysis to neurons with positive encoding did not alter our findings.

### Normalization of responses for averaging

Several figures show average traces of *offer value* response activity normalized so that values and neural responses vary in the range [0 1]. Unless otherwise specified, these responses were normalized as follows. For cases where either *V*_*max*_ or *V*_*min*_ change alone:

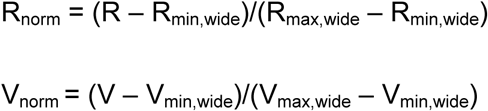

For cases where *V*_*max*_ and *V*_*min*_ shift concurrently:

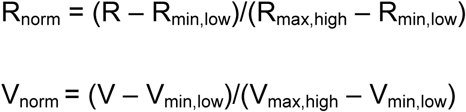

*R*_*norm*_ and *V*_*norm*_ denote normalized responses and values, *R* and *V* denote the non-normalized responses and values, and *R*_*max,j*_ and *R*_*min,j*_ indicate the response to *V*_*max*_ and *V*_*min*_ in range type *j*.

### Metrics of adaptation

Analysis of adaptation focused on *offer value A, offer value B,* and *chosen value* responses. We grouped responses into three types of range transition: change *V*_*max*_ only, change *V*_*min*_ only, and change both. Transition types could be divided further based on the direction of change (increase/decrease). For *offer value* responses, we controlled the value range so that each transition type was consistent across sessions. Thus if we describe the offer value range as a fraction of the wide value range (Δ*V*_*wide*_), the normalized ranges were 0-0.6uV (low range), 0.4-1uV (high range), and 0-1 uV (wide range) for all *offer value* responses. *Chosen value* ranges depended on the choice pattern of the animal, and in particular the relative value *ρ*, which varied across sessions even when the two juices were identical. For the purposes of this experiment, we considered the maximum/minimum chosen value changed if the difference between blocks was greater than >0.5 uB.

For each response, we regressed neural activity onto value separately in each block. We obtained the slope of encoding (*s*) from each fit. Slopes were compared directly across range types, and the relationship between slope and range was tested more precisely using Adaptation Ratios (see main text). We also used the regression to calculate the responses to the minimum and maximum values (*R*_*min*_ and *R*_*max*_) from the regression:

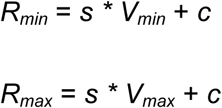

where *V*_*min*_ and *V*_*max*_ are the minimum and maximum values in the current block and *c* is the y-intercept of the linear fit. We computed the normalized difference for conditions where either *V*_*min*_ or *V*_*max*_ change alone:

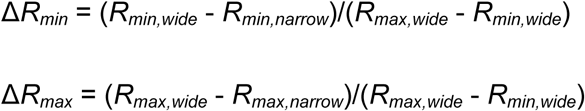

And for conditions where both change:

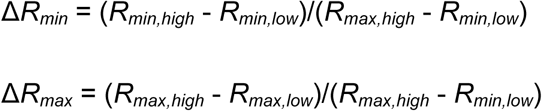

We also computed the values of Δ*R*_*min*_ and Δ*R*_*max*_ that would be predicted if neurons did not adapt at all (NA). In this case Δ*R*_*min*_ and Δ*R*_*max*_ are equivalent to the difference in *V*_*max*_ and *V*_*min*_ across conditions, normalized as above. For example, when either *V*_*max*_ or *V*_*min*_ changes alone:

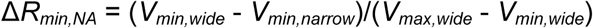

For *offer value* responses, changes in *V*_*min*_ and *V*_*max*_ are controlled. Thus, when *V*_*max*_ changes alone Δ*R*_*max,NA*_ = 0.4 and in Δ*R*_*min,NA*_ = 0; when *V*_*min*_ changes alone Δ*R*_*max,NA*_ = 0 and in Δ*R*_*min,NA*_ = 0.4; and when both change, Δ*R*_*max,NA*_ = Δ*R*_*min,NA*_ = 0.4. For *chosen value* neurons, Δ*R*_*max,NA*_ and Δ*R*_*min,NA*_ depend on the relative value and the animal’s choice pattern in each session.

### Analysis of time course

To study adaptation in early vs. late trials after the range transition, we took the first and second half of each block and computed separate tuning functions for each half. Responses were excluded if the slope changed by a factor >5 within the first block (2 responses excluded). Including these responses did not substantially affect results, but did add noise to the data, particularly for changes in Vmax. Plots of mean tuning curves in the first and second halves of each block were normalized to the first half of the wide range block.

### Simulations

We constructed a linear model of decision making to explore the effect of minimum (baseline) firing rates on choice behavior. For the purpose of the model, we defined the baseline (*R*_*min*_) as the minimum neural activity in a given block. This value could correspond to either a nonzero baseline firing rate or to the minimum evoked response in a given context. The model consisted of a population of 10,000 simulated *offer A* and *offer B* neurons (5000 units per group). Each unit encoded *offer value* in a linear way, such that the response of unit *i* on trial *t* was:

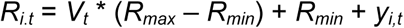

where *R*_*min*_ is the baseline activity, *R*_*max*_ is the maximum response of the unit, *V*_*t*_ is the value of the encoded juice on trial *t,* and *y*_*i,t*_ is a noise term for unit *i* on trial *t*. Units of *R* and *V* are arbitrary.

Importantly, *offer value* neurons in OFC show small but significant noise correlations (*r*_*noise*_) (Conen & Padoa-Schioppa, 2015). We generated a realistic correlation matrix Q for the population as described previously (Conen & Padoa-Schioppa, 2015; Hardin, Garcia, & Golan, 2013). We set mean(*r*_*noise*_) = 0.01 for units encoding the same juice and mean(*r*_*noise*_) = 0 for units encoding different juices. To generate the vector of noise terms y_t_ for the population on each trial, we generated values of uncorrelated noise u_t_ ∼ N(0,1). This was multiplied by the correlation matrix and scaled according to the Fano factor (*F*) and the mean response for the current offer type (⟨*R*_*V*_⟩) to obtain y_t_:

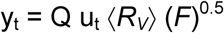

the scaling factor ⟨*R*_*V*_⟩ (*F*)^0.5^ accounts the observation that the variance in firing rate is proportional to the mean response.

Using this model, we simulated choice behavior for increasing values of *R*_*min*_. We considered two scenarios: 1) units had a fixed *R*_*max*_, or 2) units had a fixed activity range (*R*_*max*_ – *R*_*min*_). For convenience, we defined *R*_*max*_ = 1 for the first scenario and (*R*_*max*_ – *R*_*min*_) = 1 for the second. Each simulation consisted of 1000 trials, and the decision on each trial was determined by the difference in the net activity of the *offer value A* and *offer value B* units. The value of each juice for a given trial was a randomly chosen integer ranging from 0 to 10. In both scenarios, we simulated the choice pattern for the neural population as values of *R*_*min*_ increased from 0 to 1 in increments of 0.01. We repeated the process for five different values of *F* and ran the simulation 20 times for each value of *F* and *R*_*min*_. As in a previous study (Rustichini et al., 2017), we measured the effectiveness of choice behavior using fractional lost value (FLV):

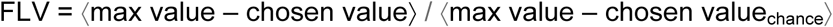

where [max value] refers to the higher value of the two offers on a given trial and [chosen value_chance_] is the average of the two offers. If a subject always chooses the max value, FLV = 0; if they choose randomly, FLV = 1.

## Acknowledgments

We thank H. Schoknecht for help with animal training and S. Ballesta, W. Shi, and E. Bromberg-Martin for comments on earlier versions of the manuscript. This work was supported by the National Institutes of Health (grant number R01-MH104494 to CPS and grant number F31-MH107111 to KEC).

## References

Adibi, M., McDonald, J. S., Clifford, C. W. G., & Arabzadeh, E. (2013). Adaptation improves neural coding efficiency despite increasing correlations in variability. Journal of Neuroscience, 33(5), 2108–2120. https://DOI.org/10.1523/JNEUROSCI.3449-12.2013

Barlow, H. B. (1961). Possible Principles Underlying the Transformations of Sensory Messages.

In W. A. Rosenblith (Ed.), Sensory Communication (pp. 217–234). Cambridge, MA: MIT Press. https://DOI.org/10.7551/mitpress/9780262518420.003.0013

Beck, J. M., Latham, P. E., & Pouget, A. (2011). Marginalization in neural circuits with divisive normalization. Journal of Neuroscience, 31(43), 15310–15319. https://DOI.org/10.1523/JNEUROSCI.1706-11.2011

Benucci, A., Saleem, A. B., & Carandini, M. (2013). Adaptation maintains population homeostasis in primary visual cortex. Nature Neuroscience, 16(6), 724–729. https://DOI.org/10.1038/nn.3382

Bermudez, M. A., & Schultz, W. (2010). Reward Magnitude Coding in Primate Amygdala Neurons. Journal of Neurophysiology, 104(6), 3424–3432. https://DOI.org/10.1152/jn.00540.2010

Burke, C. J., Baddeley, M., Tobler, P. N., & Schultz, W. (2016). Partial adaptation of obtained and observed value signals preserves information about gains and losses. Journal of Neuroscience, 36(39), 10016–10025. https://DOI.org/10.1523/JNEUROSCI.0487-16.2016

Cai, X., & Padoa-Schioppa, C. (2014). Contributions of orbitofrontal and lateral prefrontal cortices to economic choice and the good-to-action transformation. Neuron, 81(5), 1140–1151. https://DOI.org/10.1016/j.neuron.2014.01.008

Carandini, M., & Heeger, D. J. (2011). Normalization as a canonical neural computation. Nature Reviews Neuroscience, 13(1), 51–62. https://DOI.org/10.1038/nrn3136

Chance, F. S., Abbott, L. F., & Reyes, A. D. (2002). Gain modulation from background synaptic input. Neuron, 35(4), 773–782. https://DOI.org/10.1016/S0896-6273(02)00820-6

Conen, K. E., & Padoa-Schioppa, C. (2015). Neuronal variability in orbitofrontal cortex during economic decisions. Journal of Neurophysiology, 114(3), 1367–1381. https://DOI.org/10.1152/jn.00231.2015

Cox, K. M., & Kable, J. W. (2014). BOLD subjective value signals exhibit robust range adaptation. Journal of Neuroscience, 34(49), 16533–16543. https://DOI.org/10.1523/JNEUROSCI.3927-14.2014

Dan, Y., Atick, J. J., & Reid, R. C. (1996). Efficient coding of natural scenes in the lateral geniculate nucleus: experimental test of a computational theory. Journal of Neuroscience, 16(10), 3351–3362.

Díaz-Quesada, M., & Maravall, M. (2008). Intrinsic mechanisms for adaptive gain rescaling in barrel cortex. Journal of Neuroscience, 28(3), 696–710. https://DOI.org/10.1523/JNEUROSCI.4931-07.2008

Elliott, R., Agnew, Z., & Deakin, J. F. W. (2008). Medial orbitofrontal cortex codes relative rather than absolute value of financial rewards in humans. European Journal of Neuroscience, 27(9), 2213–2218. https://DOI.org/10.1111/j.1460-9568.2008.06202.x

Fairhall, A. L., Lewen, G. D., Bialek, W., & De Ruyter van Steveninck, R. R. (2001). Efficiency and ambiguity in an adaptive neural code. Nature, 412(6849), 787–792. https://DOI.org/10.1038/35090500

Fellows, L. K. (2011). Orbitofrontal contributions to value-based decision making: Evidence from humans with frontal lobe damage. Annals of the New York Academy of Sciences, 1239(1), 51–58. https://DOI.org/10.1111/j.1749-6632.2011.06229.x

Gutnisky, D. A., & Dragoi, V. (2008). Adaptive coding of visual information in neural populations. Nature, 452(7184), 220–224. https://DOI.org/10.1038/nature06563

Hardin, J., Garcia, S. R., & Golan, D. (2013). A method for generating realistic correlation matrices. Annals of Applied Statistics, 7(3), 1733–1762. https://DOI.org/10.1214/13- AOAS638

Hengen, K. B., Lambo, M. E., Van Hooser, S. D., Katz, D. B., & Turrigiano, G. G. (2013). Firing rate homeostasis in visual cortex of freely behaving rodents. Neuron, 80(2), 335–342. https://DOI.org/10.1016/J.NEURON.2013.08.038

Higgs, M. H. (2006). Diversity of gain modulation by noise in neocortical neurons: regulation by the slow afterhyperpolarization conductance. Journal of Neuroscience, 26(34), 8787–8799. https://DOI.org/10.1523/JNEUROSCI.1792-06.2006

Holt, G. R., & Koch, C. (1997). Shunting inhibition does not have a divisive effect on firing rates. Neural Computation, 9(5), 1001–1013. https://DOI.org/10.1162/neco.1997.9.5.1001

Kobayashi, S., Pinto de Carvalho, O., & Schultz, W. (2010). Adaptation of reward sensitivity in orbitofrontal neurons. Journal of Neuroscience, 30(2), 534–544. https://DOI.org/10.1523/JNEUROSCI.4009-09.2010

Kohn, A. (2007). Visual adaptation: physiology, mechanisms, and functional benefits. Journal of Neurophysiology, 97, 3155–3164. https://DOI.org/10.1152/jn.00086.2007

Krekelberg, B., van Wezel, R. J. A., & Albright, T. D. (2006). Adaptation in macaque MT reduces perceived speed and improves speed discrimination. Journal of Neurophysiology, 95(1), 255–270. https://DOI.org/10.1152/jn.00750.2005

Laughlin, S. (1981). A simple coding procedure enhances a neuron’s information capacity. Zeitschrift Fur Naturforschung C, 36, 910–912. https://DOI.org/10.1515/znc-1981-9-1040

Lewicki, M. S. (2002). Efficient coding of natural sounds. Nature Neuroscience, 5(4), 356–363. https://DOI.org/10.1038/nn831

Liu, B., Macellaio, M. V., & Osborne, L. C. (2016). Efficient sensory cortical coding optimizes pursuit eye movements. Nature Communications, 7, 12759. https://DOI.org/10.1038/ncomms12759

Mease, R. A., Famulare, M., Gjorgjieva, J., Moody, W. J., & Fairhall, A. L. (2013). Emergence of adaptive computation by single neurons in the developing cortex. Journal of Neuroscience, 33(30), 12154–12170. https://DOI.org/10.1523/JNEUROSCI.3263-12.2013

Natan, R. G., Rao, W., & Geffen, M. N. (2017). Cortical interneurons differentially shape frequency tuning following adaptation. Cell Reports, 21(4), 878–890. https://DOI.org/10.1016/j.celrep.2017.10.012

Ohshiro, T., Angelaki, D. E., & DeAngelis, G. C. (2011). A normalization model of multisensory integration. Nature Neuroscience, 14(6), 775–782. https://DOI.org/10.1038/nn.2815

Olsen, S. R., Bhandawat, V., & Wilson, R. I. (2010). Divisive normalization in olfactory population codes. Neuron, 66(2), 287–299. https://DOI.org/10.1016/J.NEURON.2010.04.009

Olsen, S. R., Bortone, D. S., Adesnik, H., & Scanziani, M. (2012). Gain control by layer six in cortical circuits of vision. Nature, 483(7387), 47–52. https://DOI.org/10.1038/nature10835

Ongur, D., & Price, J. (2000). The organization of networks within the orbital and medial prefrontal cortex of rats, monkeys and humans. Cerebral Cortex, 10(3), 206–219. https://DOI.org/10.1093/cercor/10.3.206

Padoa-Schioppa, C. (2009). Range-adapting representation of economic value in the orbitofrontal cortex. Journal of Neuroscience, 29(44), 1404–14014. https://DOI.org/10.1523/JNEUROSCI.3751-09.2009

Padoa-Schioppa, C., & Assad, J. A. (2006). Neurons in the orbitofrontal cortex encode economic value. Nature, 441(7090), 223–226. https://DOI.org/10.1038/nature04676

Padoa-Schioppa, C., & Conen, K. E. (2017). Orbitofrontal cortex: a neural circuit for economic decisions. Neuron. https://DOI.org/10.1016/j.neuron.2017.09.031

Rudebeck, P. H., & Murray, E. A. (2014). The orbitofrontal oracle: cortical mechanisms for the prediction and evaluation of specific behavioral outcomes. Neuron, 84(6), 1143–1156. https://DOI.org/10.1016/j.neuron.2014.10.049

Rustichini, A., Conen, K. E., Cai, X., & Padoa-Schioppa, C. (2017). Optimal coding and neuronal adaptation in economic decisions. Nature Communications, 8(1). https://DOI.org/10.1038/s41467-017-01373-y

Saez, R. A., Saez, A., Paton, J. J., Lau, B., & Salzman, C. D. (2017). Distinct roles for the amygdala and orbitofrontal cortex in representing the relative amount of expected reward. Neuron, 95(1), 70–77.e3. https://DOI.org/10.1016/j.neuron.2017.06.012

Sanchez-Vives, M. V, Nowak, L. G., & McCormick, D. A. (2000). Cellular mechanisms of long-lasting adaptation in visual cortical neurons in vitro. Journal of Neuroscience, 20(11), 4286–4299. https://DOI.org/20/11/4286 [pii]

Sanchez-Vives, M. V, Nowak, L. G., & McCormick, D. A. (2000). Membrane mechanisms underlying contrast adaptation in cat area 17 in vivo. Journal of Neuroscience, 20(11), 4267–4285. https://DOI.org/20/11/4267 [pii]

Schultz, W. (2015). Neuronal reward and decision signals: from theories to data. Physiological Reviews, 95(3), 853–951. https://DOI.org/10.1152/physrev.00023.2014

Simoncelli, E. P., & Schwartz, O. (2001). Natural sound statistics and divisive normalization in the auditory system. Advances in Neural Information Processing Systems, 13, 27–30.

Soltani, A., De Martino, B., & Camerer, C. (2012). A range-normalization model of context-dependent choice: a new model and evidence. PLoS Computational Biology, 8(7), e1002607. https://DOI.org/10.1371/journal.pcbi.1002607

Valerio, R., & Navarro, R. (2003). Optimal coding through divisive normalization models of V1 neurons. In Network: Computation in Neural Systems (Vol. 14, pp. 579–593). https://DOI.org/10.1088/0954-898X/14/3/310

Varela, J. A., Sen, K., Gibson, J., Fost, J., Abbott, L. F., & Nelson, S. B. (1997). A quantitative description of short-term plasticity at excitatory synapses in layer 2/3 of rat primary visual cortex. Journal of Neuroscience, 17(20), 7926–7940. https://DOI.org/10.1523/JNEUROSCI.17-20-07926.1997

Wallis, J. D. (2012). Cross-species studies of orbitofrontal cortex and value-based decision-making. Nature Neuroscience, 15(1), 13–19. https://DOI.org/10.1038/nn.2956

Wark, B., Lundstrom, B. N., & Fairhall, A. (2007). Sensory adaptation. Current Opinion in Neurobiology, 17(4), 423–429. https://DOI.org/10.1016/j.conb.2007.07.001

Wilson, N. R., Runyan, C. A., Wang, F. L., & Sur, M. (2012). Division and subtraction by distinct cortical inhibitory networks in vivo. Nature, 488(7411), 343–348. https://DOI.org/10.1038/nature11347

Yamada, H., Louie, K., Tymula, A., & Glimcher, P. W. (2018). Free choice shapes normalized value signals in medial orbitofrontal cortex. Nature Communications, 9(1). https://DOI.org/10.1038/s41467-017-02614-w

Zimmerman, J., Glimcher, P., & Louie, K. (2018). Multiple timescales of normalized value coding underlie adaptive choice behavior. Nature Communications, 9, 3206. https://DOI.org/10.1038/s41467-018-05507-8

